# A Bayesian Analysis of Steady–State Enzyme Data leads to Estimates of Rate Constants and Uncertainties in a Multi-Step Reaction, along with Free Energy Profiles^†^

**DOI:** 10.1101/2021.08.04.454956

**Authors:** Ian Barr

## Abstract

The microscopic rate constants that govern an enzymatic reaction are only directly measured under certain experimental set-ups, such as stopped flow, quenched flow, or temperature-jump assays; the majority of enzymology proceeds from steady state conditions which leads to a set of more easily–observable parameters such as *k*_*cat*_, *K*_*M*_, and observed Kinetic Isotope Effects 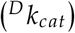. This paper further develops a model from Toney (2013) to estimate microscopic rate constants from steady-state data for a set of reversible, four–step reactions. This paper uses the Bayesian modeling software Stan, and demonstrates the benefits of Bayesian data analysis in the estimation of these rate constants. In contrast to the optimization methods employed often in the estimation of kinetic constants, a Bayesian treatment is more equipped to estimate the uncertainties of each parameter; sampling from the posterior distribution using Hamiltonian Monte Carlo immediately gives parameter estimates as mean or median of the posterior, and also confidence intervals that express the uncertainty of each parameter.

## 1 Introduction

Estimation of the rate constants associated with each step of an enzymatic mechanisms is rarely straightforward, due to complexity of the reactions and lack of an ability to observe each intermediate species during the course of a reaction. The two enzymes under study here are alanine racemase (AR, EC 5.1.1.1), which catalyzes the reversible conversion of l-alanine to d-alanine, and triosephosphate isomerase (TIM, EC 5.3.1.1), which functions in glycolysis to convert dihydroxyacetone phosphate into d-glyceraldehyde 3-phosphate. Both are classified as isomerases, and take a single substate in both the forward and reverse directions. The general reaction scheme for AR and TIM is given in Scheme 1. In order to fully characterize these reactions, kinetically,

**Scheme 1:**
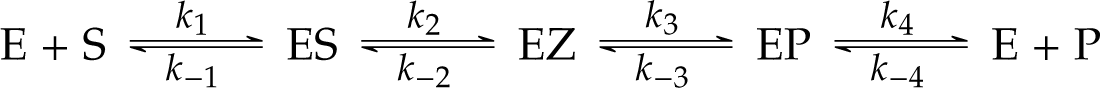
The Reaction scheme for a reversible, 4-step reaction. ^2^

we would like to estimate the rate constants for every step. In addition, if certain rate constants are isotopically sensitive, there will be additional values to estimate. For an enzymatic reaction scheme with four reversible steps, that leaves us with 8 microscopic rate constants to determine. In Scheme 1, *k*_1_ and *k*_−4_ are second–order rate constants, and all others are first order. EZ is an intermediate that reacts rapidly in both directions. The substrates are taken to be l-alanine for AR and dihydroxyacetone phosphate for TIM, though the reactions are reversible. First order rate constants *k*_2_ and *k*_−3_ are isotopically sensitive, with primary kinetic isotope effects 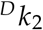 and 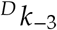.

Since we cannot directly measure *k*_1_, *k*_2_, etc., we have to rely on indirect methods of determining those values. Ref. 1, which is the starting point for this work, uses a series of measurements done under steady state condition, each of which can be related to the microscopic rate constants mathematically (Eqs. (2)–(12)). Through incorporation of sufficient experimental data, it is possible in principle to determine each of the microscopic rate constants. In Ref. 1, global fitting is used to extract individual rate constants from steady-state reaction data. Global fitting in this case refers to the use of a target function containing contributions from all of the experimental data, from which are estimated a set of parameters consistent with the entire data set through non-linear regression. The earlier work used standard non-linear optimization algorithms to minimize the relative squared error of a set of data points. The target function used was

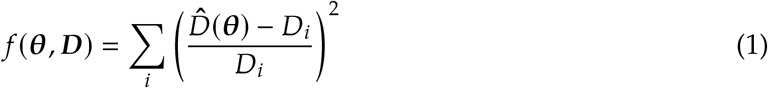

Where θ is a vector of parameters to be estimated, *D* is a vector of experimental values, and 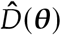 is the function relating the parameters to the experimental value. This function leads to a minimization of the relative standard deviation (RSD), which is preferred because the experimental values are different orders of magnitude so they must be scaled to avoid bias. Ref. 1 showed that convergence was achievable using non-linear optimization, and that the method was reasonably robust. The fact that an optimization algorithm converges on a set of parameter values is not in itself useful, unless we have some confidence in those numbers. Ref. 1 wisely uses a method whereby a set of randomly generated values with the same mean and standard deviation as the experimental data are fed into the optimization algorithm, and parameters are re-calculated for each set, allowing an estimation of parameter uncertainty. Other non-linear methods would employ the Hessian matrix or bootstrapping to the same effect. ^3,4^ These methods fall under the rubric of frequentist analysis, which is often faster and equally as accurate as Bayesian methods are, given plentiful, high quality data. However, when the number of parameters to be estimated is nearly equal to the number of data points, as in the current case, Bayesian methods can provide invaluable information about the most likely parameter values, given all available data, and the uncertainty the estimates of each parameter. ^5^ Here I show that a Bayesian modeling of the same system gives robust and useful estimates of the rate constants and their associated uncertainties. In addition, a Bayesian treatment is able to handle cases of possible experimental error, at the cost of greater uncertainty in the parameter posterior distributions. ^6^

### 1.1 Incorporation of the Equilibrium Constant

We have here introduced a new data value to improve the estimate, the equilibrium constant *K*_*eq*_ (Eq. 13). This is equal to the product of the forward rate constants divided by the reverse rate constants, and can be determined experimentally by measuring the concentrations of reactant and product at equilibrium, or indirectly from the forward and reverse *k*_*cat*_/*K*_*m*_ values using the Haldane relationship: ^11,12^

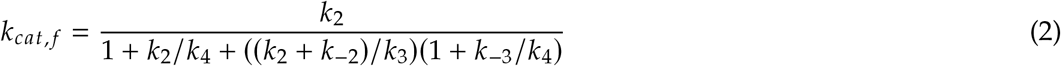

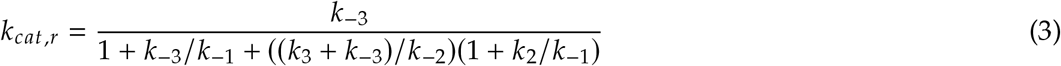

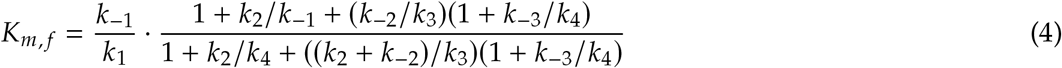

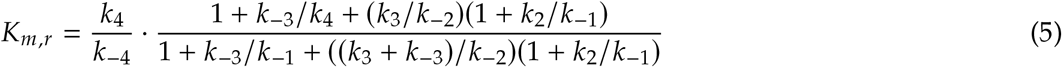

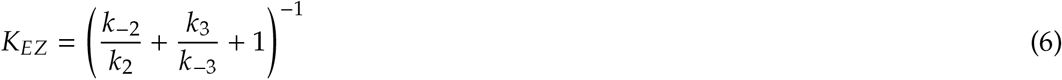

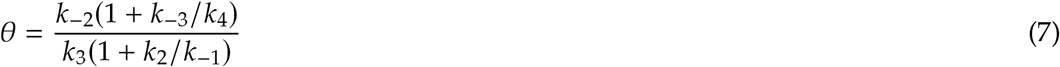

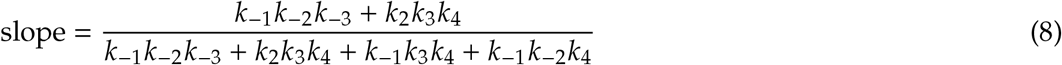

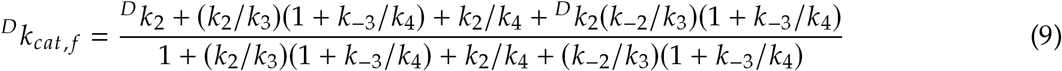

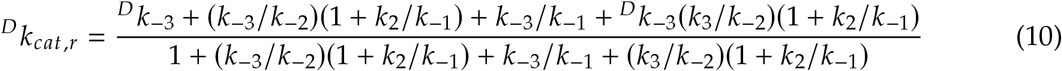

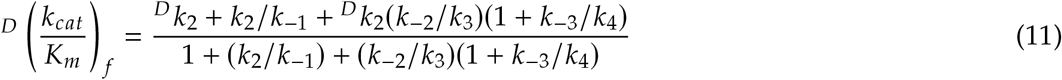

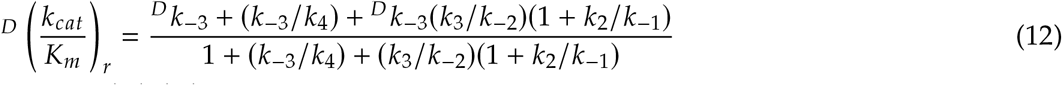

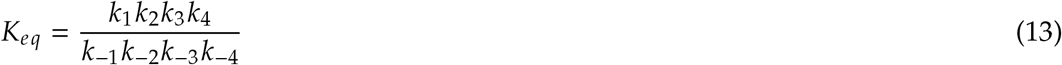

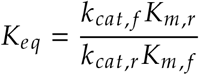

Direct measurement of *K*_*eq*_ is to be preferred, since use of the Haldane relationship utilizes the *k*_*cat*_ and *K*_*m*_ values that are already incorporated into the model, so using these values again tends to bias the estimates. For the same reason, the values of (^*D*^ (*k*_*cat*_/*K*_*m*_) − 1 / ^*D*^ *k*_*cat*_ − 1)used in Ref. 1 are not used here, because they represented re-use of data that is already incorporated as ^*D*^ *k*_*cat*_ and ^*D*^ (*k*_*cat*_/*K*_*m*_). However, in some cases the *K*_*eq*_ might be hard to measure directly, and the Haldane relationship may be used (with caution). More reliable estimates for *K*_*eq*_ might be possible using the Haldane relationship if there exists high quality data for homologues of the enzyme, or point mutants, because the *K*_*eq*_ values calculated by the Haldane relationship should theoretically be the same for all active versions of an enzyme, as long as the temperature and buffer composition are similar. In this case we can average the values from several sources to obtain a more reliable estimate for *K*_*eq*_.

An additional reason for using the value of *K*_*eq*_ is that the expression contains *k*_1_ and *k*_−4_, which each only appear in one other equation (for *K*_*m*_, forward and reverse). This means that we are dependent on accurate measurement of *K*_*m*_ to get reliable values for *k*_1_ and *k*_−4_, in the absence of any further information. For enzymes such as AR, which converts l-alanine to d-alanine, the *K*_*eq*_ is theoretically exactly 1, since there is no reason l-alanine would have a higher or lower free energy than d-alanine in a mostly achiral aqueous solution. For TIM, there is no direct measurement of the *K*_*eq*_ available in the literature, possibly because both dihydroxyacetone phosphate and d-glyceraldehyde 3-phosphate are themselves in equilibrium with their catalytically-inactive hydrated forms. ^8^ So in order to obtain the *K*_*eq*_ for the unhydrated forms, I averaged 4 literature values for *K*_*eq*_, derived from the Haldane relationship. ^8–10^

## 2. Considerations for Accurate Parameter estimation

### 2.1 General Limitation of the Bayesian Method

Every form of parameter estimation rests on a set of assumptions about the data and a model; this case is no different. Stan, as with other Bayesian modeling software, requires these assumptions be made explicit. Each parameter needs a prior distribution, which can affect the final result. The form of the model partly determines the results, and an incorrect model will lead to unhelpful results.

### 2.2 Choice of Priors for *k*s

One aspect of a Bayesian analysis that differs from the function minimization procedures used in Ref. 1 is the requirement to specify a prior distribution for each of the parameters. This is information that is incorporated into the model according to the modified version of Bayes’ Law: ^13^

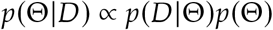

Here, the posterior distribution of the parameters *p*(Θ|*D*), the output of our simulation, is the product of the likelihood function *p*(*D* |Θ) and the prior distribution for the parameters *p*(Θ). I have chosen a uninformative prior *k* ∼ Exponential for each of the *k*s, based on the following assumptions:

1. The value of *k* is necessarily *>* 0, so an exponential distribution has the same domain.

2. The exponential distribution is often seen in physically relevant phenomena. ^14,15^

3. Setting *k* ∼ Exponential(β) with β *<<* 1 gives a broad distribution that covers the region from 1 to 1 × 10^9^, typical values for microscopic rate constants.

4. Nonetheless, the prior is not too restrictive, because we have poor prior information about which values are typical for a rate constant.

This last point is especially important, as too restrictive a prior can end up determining the shape of the posterior distribution in the absence of sufficient experimental data.

The prior for *k*_2_ (and the other *k*s) is implemented as follows in Stan:

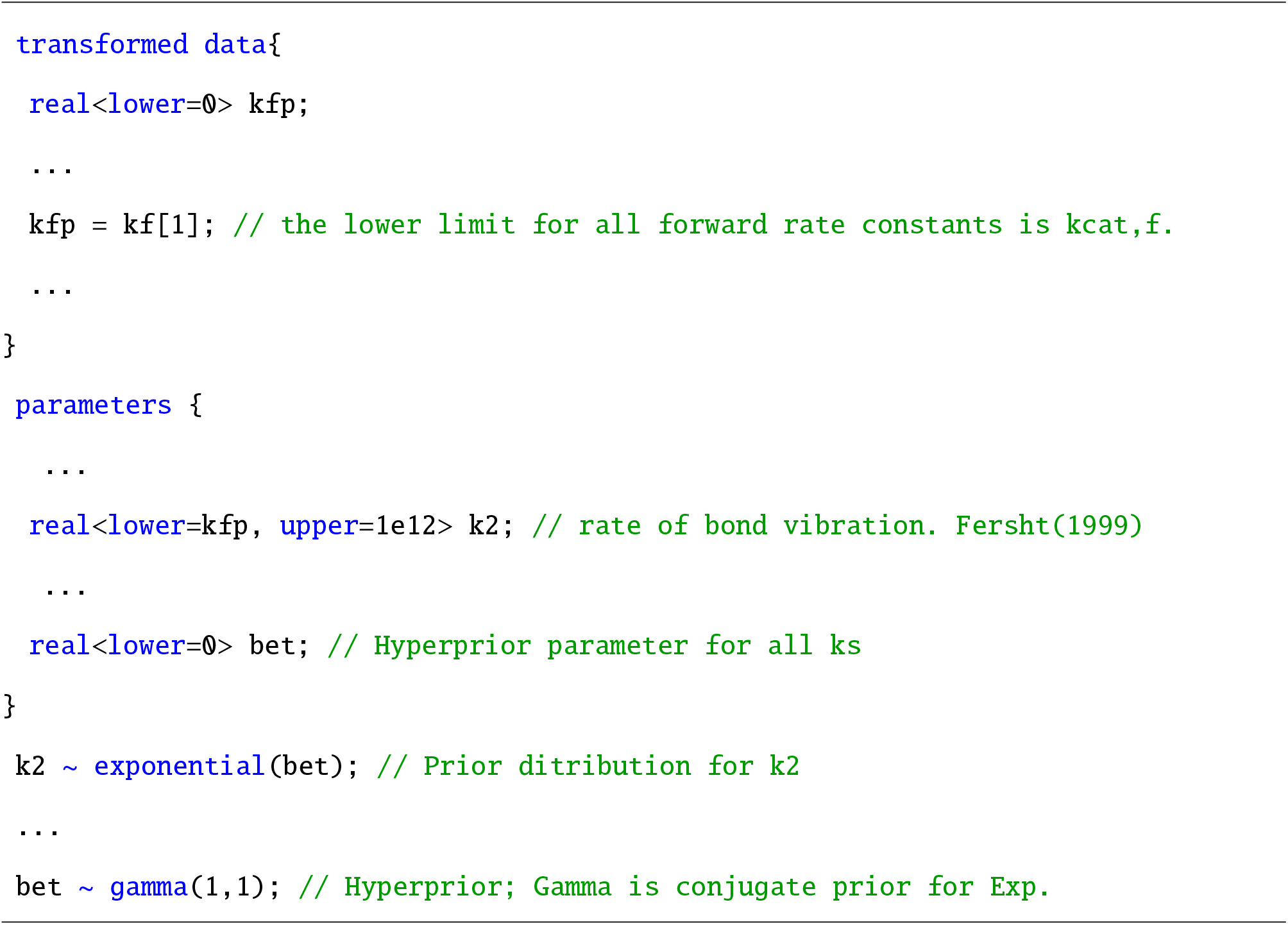

Here, we utilize a hyperprior β; the prior distribution for *k*_2_ depends on the parameter β, which is also estimated over the course of the simulation. This allows a great deal of flexibility while keeping the mathematical form of the priors constant. The hyperprior for βis set as β∼ Gamma (1, 1), a relatively uninformative prior with most of the mass below 1.

### 2.3 Choice of Priors for Intrinsic KIEs Dk_i_

Kinetic isotope effects are strictly positive quantities, and for the comparison between deuterium and protium the intrinsic KIE of step *i* is

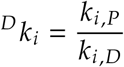

where *k*_*i,P*_ and *k*_*i,D*_ are the rate constants of the reaction with protonated and deuterated substrate. Common ranges for primary KIEs are 1.5 – 3, in the absence of quantum-mechanical tunneling. ^16^ Rarely, inverse KIEs are observed where ^*D*^ *k*_*i*_ *<* 1. Given these constraints, I set the prior as

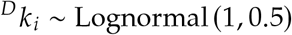

Figure 10 graphs the prior used for KIEs. We see that most of the mass is between 1 and 4, but the density extends to infinity in the positive direction. I limit the value of KIEs to less than 500, based on the fact that the largest measured enzymatic ^*D*^ *k*_*cat*_ is around 500. ^17^ Any KIE greater than 6 is likely to be due to quantum mechanical effects, and in cases where this is suspected (e.g. hydride transfer) the prior could be adjusted to reflect the expected ranges of values.

### 2.4 The Problem constants – *k*_1_ and *k*_−4_

In Ref. 1 and here, there are difficulties in accurately determining *k*_1_ and *k*_−4_ for both TIM and AR. Significantly, in Ref. 1 *k*_1_ and *k*_−4_ each only appear in one equation, the one for *K*_*m, f*_ (Eq. 4) and *K*_*m, f*_ (Eq. 5).

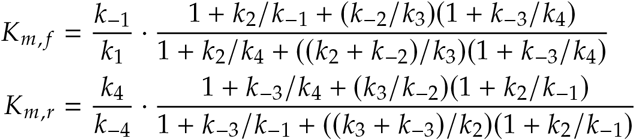

The intuitive effect of this is that each of the experimental values *besides K*_*m, f*_ and *K*_*m,r*_ only indirectly provide information as to the true value of *k*_1_ and *k*_−4_, by helping to determine the values of the other parameters. But an interesting effect of this can be seen in Figure 7, which shows correlation between parameters during the course of the simulation as the posterior distribution is explored.

**Figure 1:** Equations used in the analysis of data for TIM, AR and the simulated data. Eqs. (2)–(12) are used in Ref. 1, and Eq. (13) is included in this analysis.

**Figure 2:**
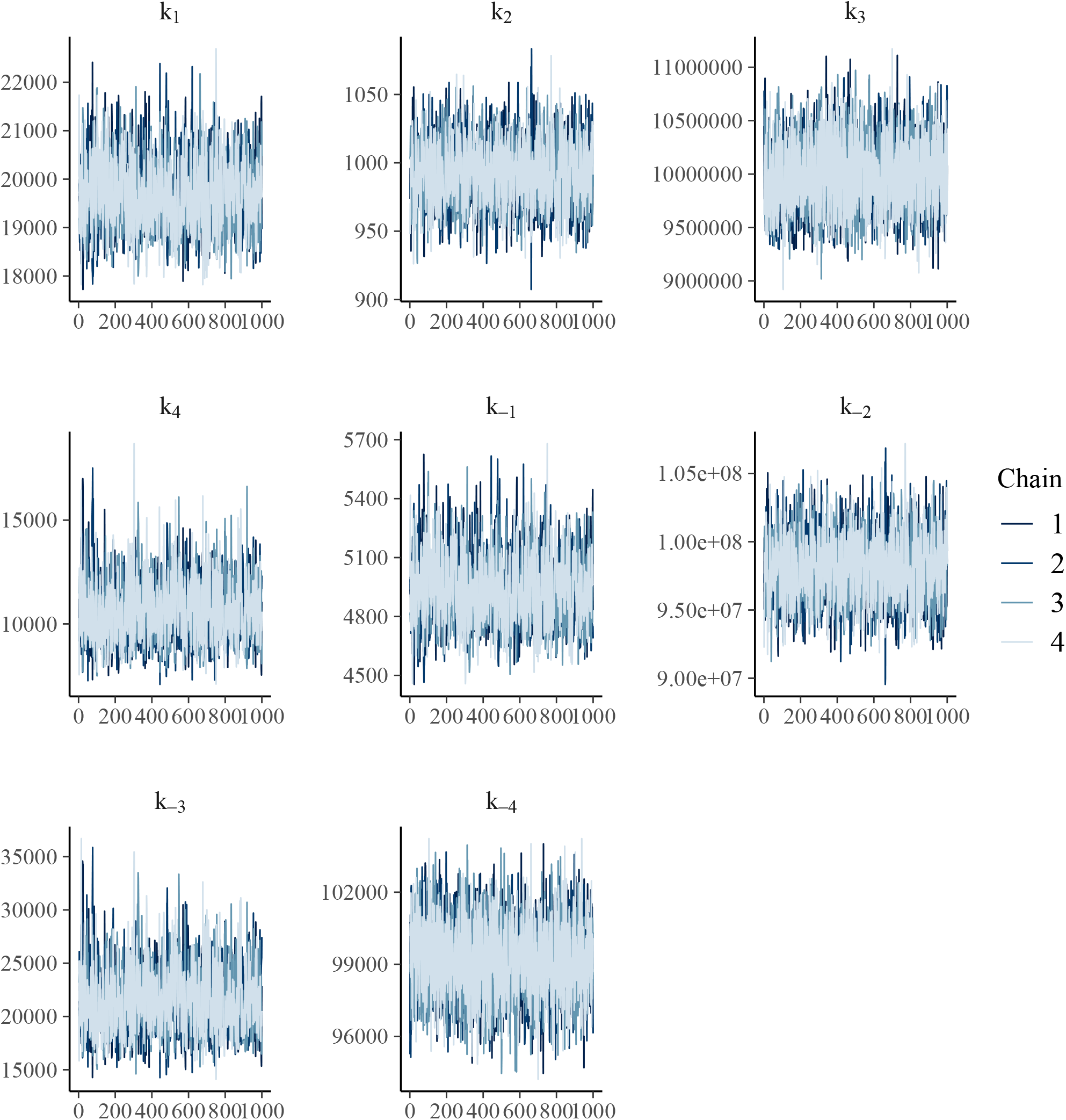
Traceplots of the modeled rate constants for the simulated data set, with experimental SD set to 0.01 for all quantities. The *y*-axis is the parameter value at each draw, and the *x*-axis are the sample numbers, post warm-up.

**Figure 3:**
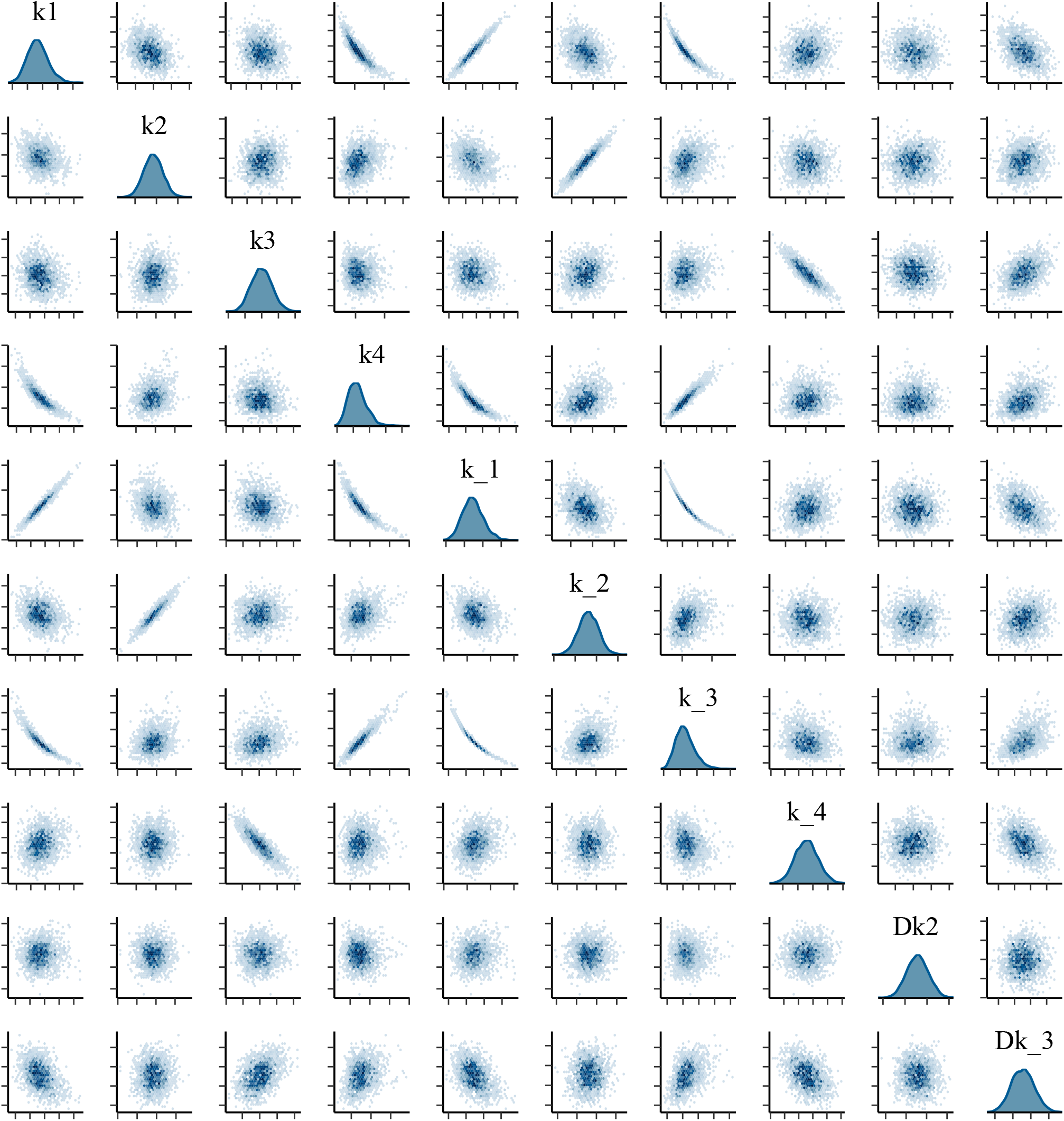
Pairwise comparison of the MCMC Draws for simulated rate constants, showing correlations between parameters.

**Figure 4:**
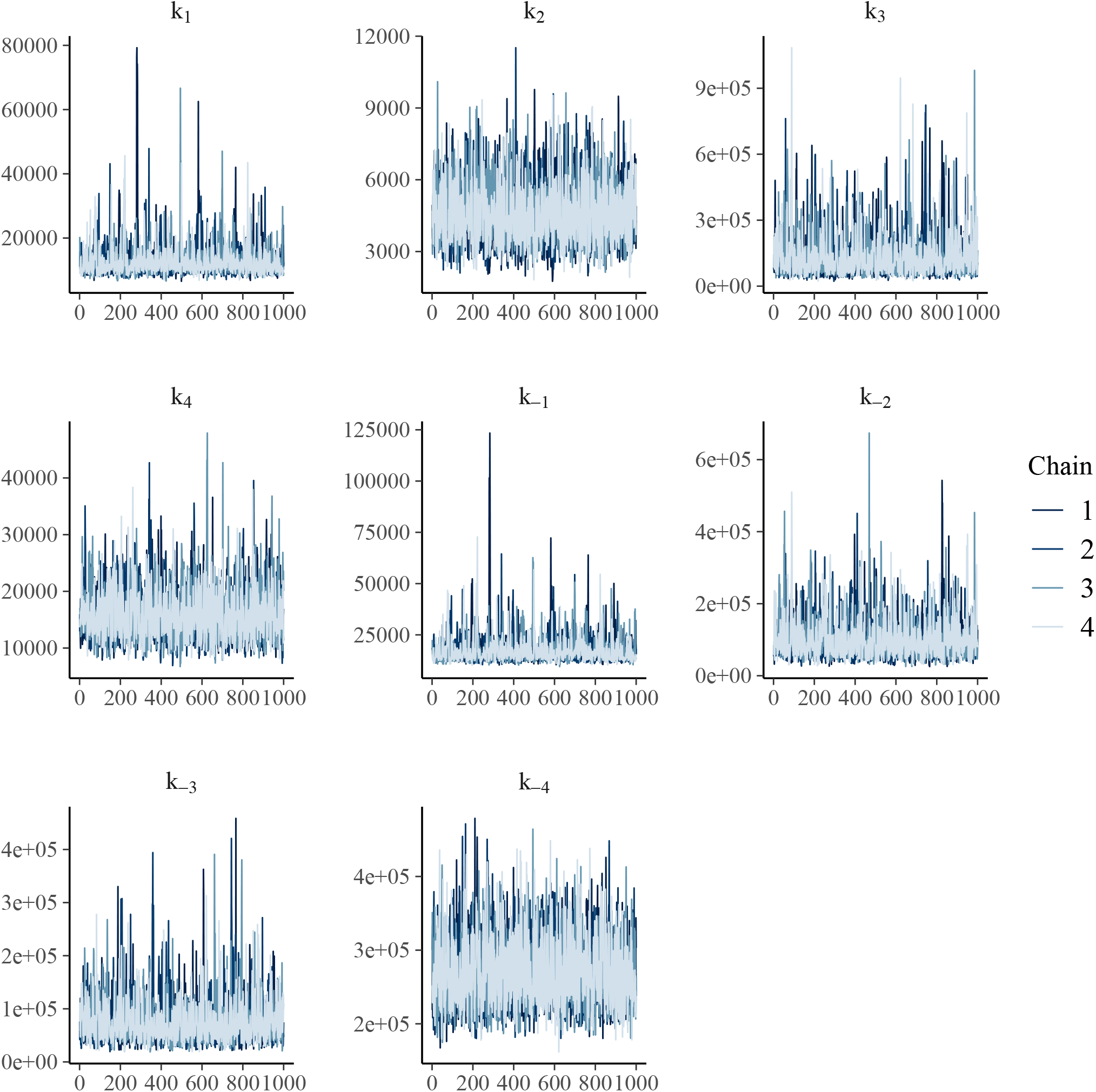
Traceplots of the rate constants for TIM.

**Figure 5:**
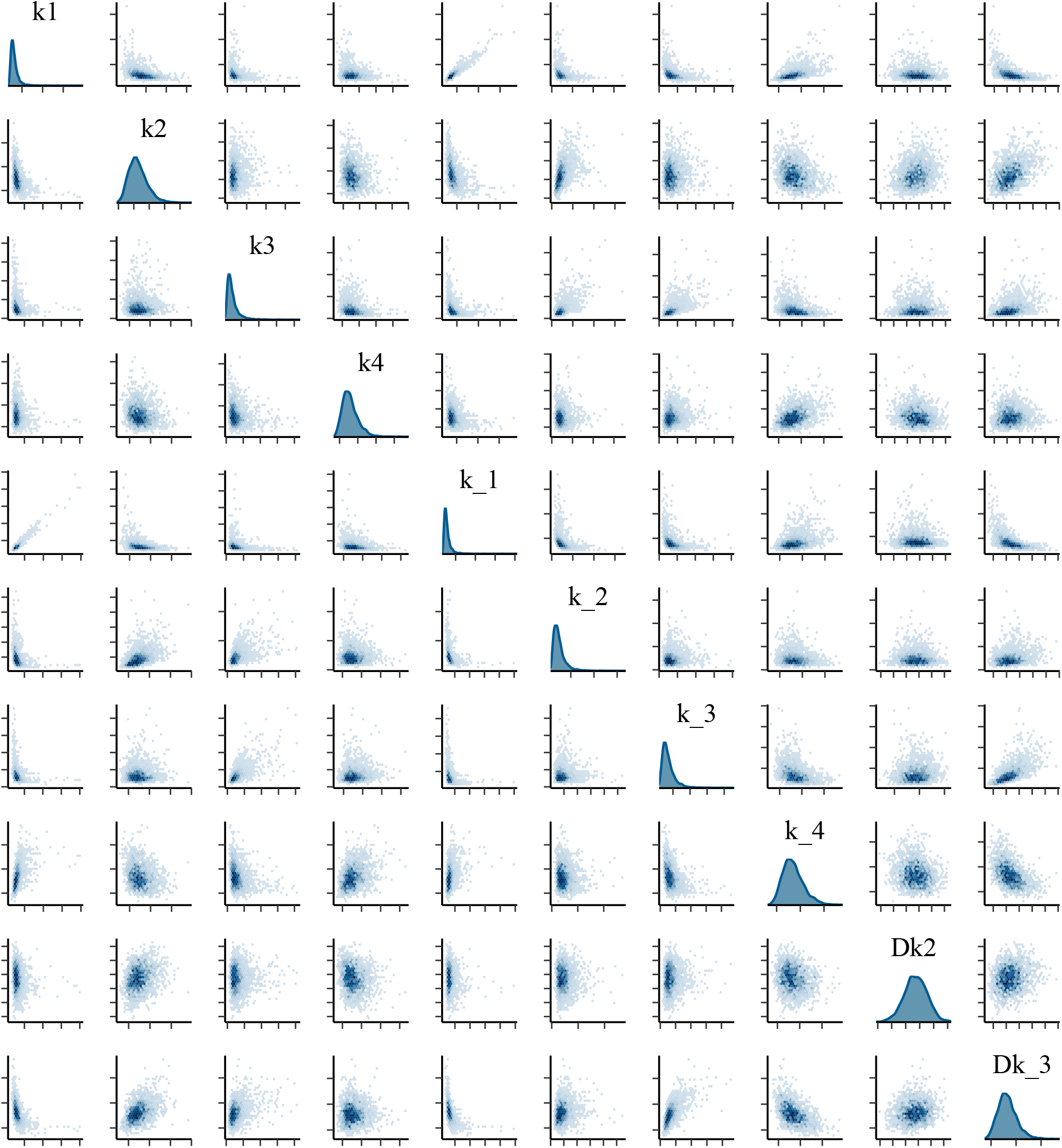
Pairwise comparison of the MCMC Draws for TIM rate constants, showing correlation between *k*_1_ and *k*_−1_

**Figure 6:**
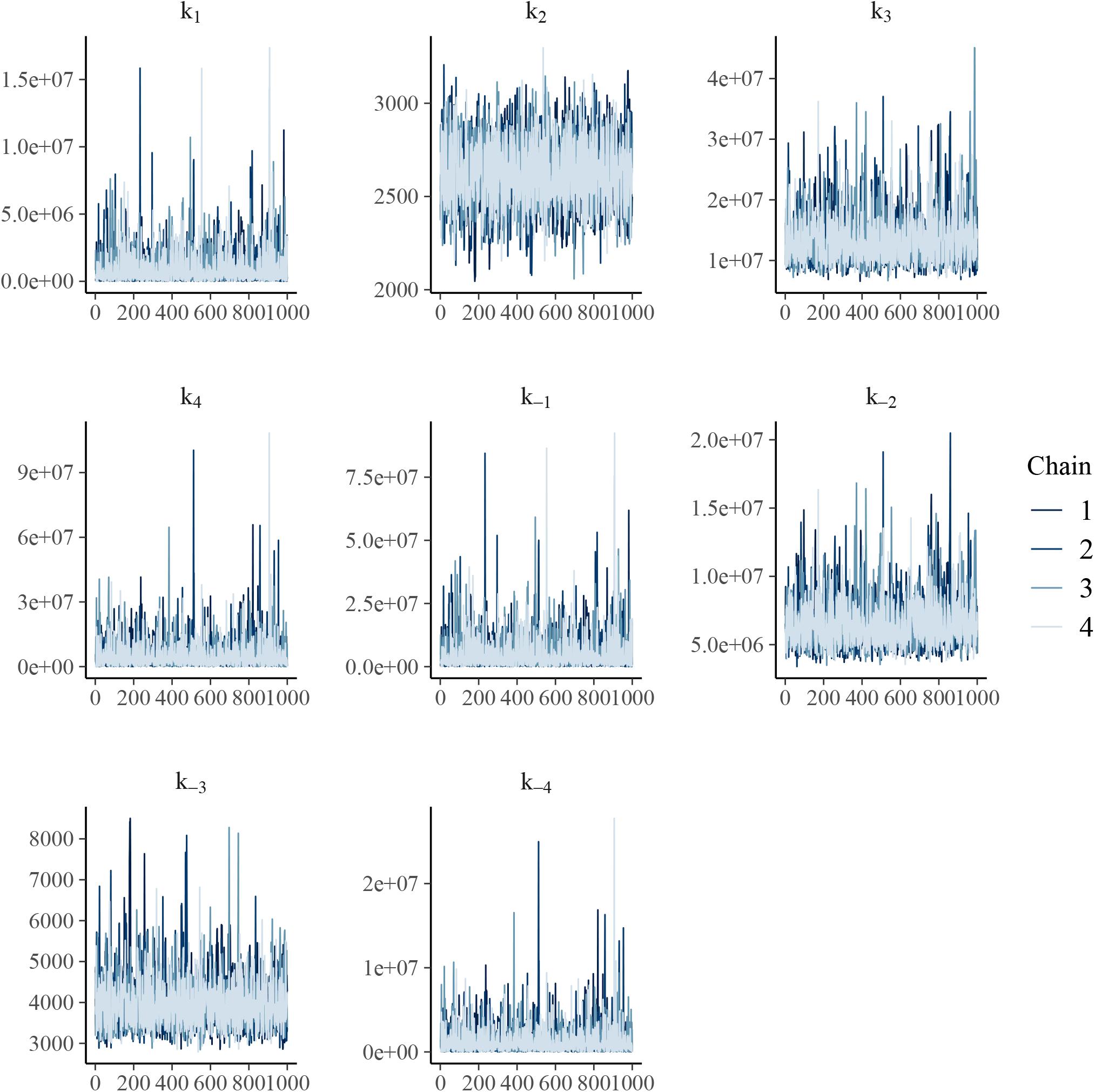
Traceplots of the rate constants for AR.

**Figure 7:**
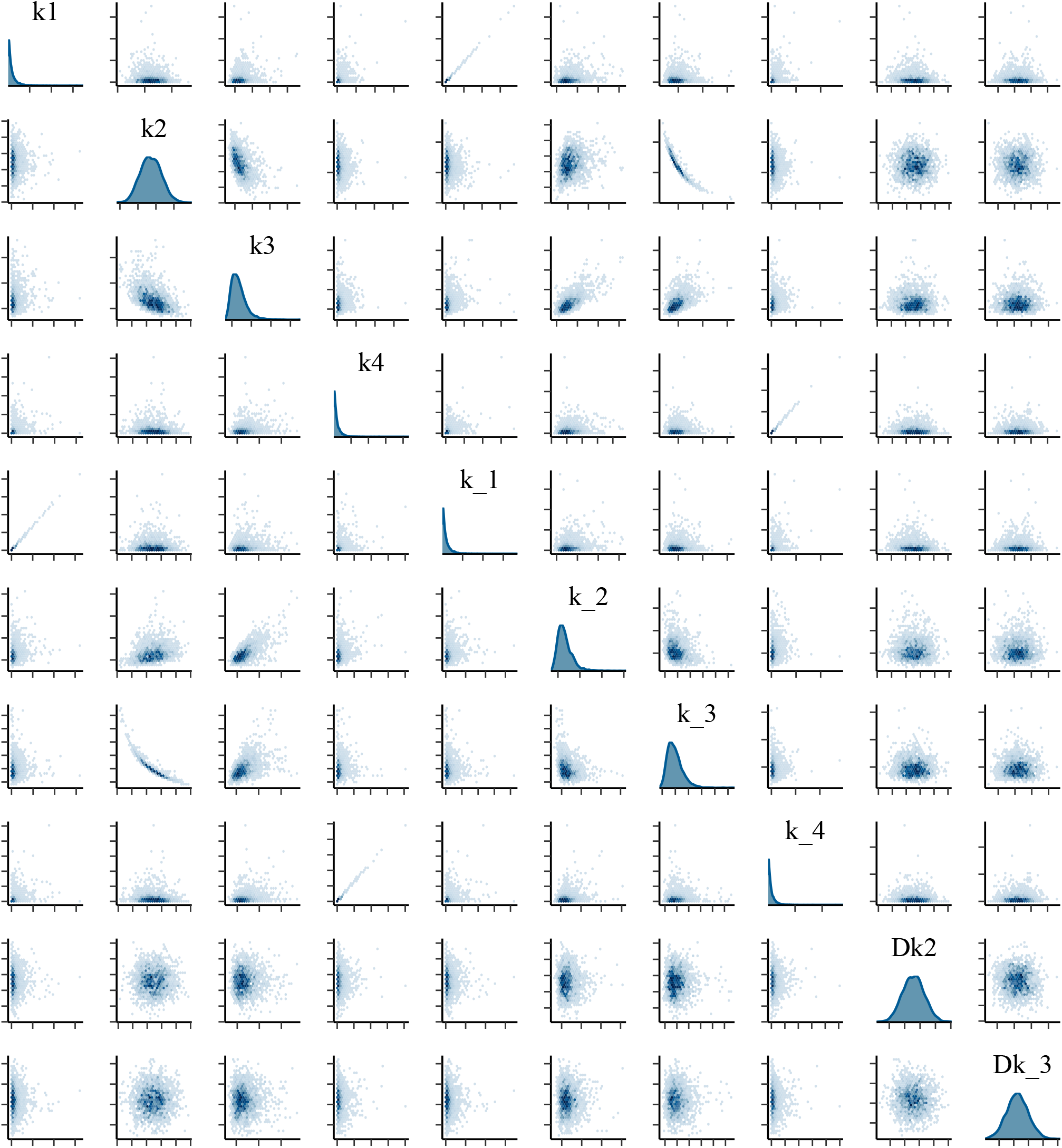
Pairwise comparison of the MCMC Draws for AR rate constants, showing correlation between *k*_1_ and *k*_−1_

In row 5, column 1, we see that the values of *k*_1_ and *k*_−1_ are linearly correlated, as are the values of *k*_4_ and *k*_−4_ in row 8, column 4. Looking at the equations for *K*_*m*_, we see that this is largely due to the fact that each contains the factor *k*_1_/*k*_−1_ or *k*_4_/*k*_−4_, and since this is the sole place that *k*_1_ and *k*_−4_ appear in this model, ambiguity in *k*_−1_ is passed along to *k*_1_, etc. Adding the data for the *K*_*eq*_ doesn’t alter this, as the expression for *K*_*eq*_ also contains *k*_1_/*k*_−1_ and *k*_4_/*k*_−4_. This tells us that *k*_1_ and *k*_−4_ can’t be considered separately from *k*_−1_ and *k*_4_; all this model can give us, in the absence of strong prior information about *k*_1_ and *k*_−4_, is the ratios *k*_1_/*k*_−1_ and *k*_4_/*k*_−4_, i.e. the equilibrium constants for the first and fourth steps. Thus in my Stan code I have replaced *k*_1_/*k*_−1_ and *k*_4_/*k*_−4_, where they appear, with *K*_1_ and *K*_4_. This slightly simplifies the calculations, and for a reversible reaction such as these it is reasonable to assume that the forward and reverse constants are within three orders of magnitude of each other, so we can limit the value of *K*_1_ and *K*_4_ during the simulation to between 0 and 10^3^. Indeed, in both TIM and AR the values determined are approximately equal to *K*_*m, f*_ and 1/*K*_*m,r*_, though this is not necessarily true in general as *K*_*m*_s can be greater than, less than, or equal to the association equilibrium constant (e.g. *K*_1_) in the case of a multi-step reaction. ^18^

## 3. Results and discussion

### 3.1 Application to Simulated Data

We base our estimates on a set of 12 equations, and we estimate 11 parameters from these data points and their uncertainties, following the general rule of thumb that one can estimate at best *n* − 1 unknown parameters from *n* data points. However, this is only best–case; experimental error and the structure of the model can limit our ability to estimate parameters effectively. The primary difficulty here is one of structural identifiability; ^19^ can we, even with ideal data, estimate the parameters given the model we have?

To test the ability of our model to accurately determine rate constants, I simulated a data set with a fixed relative standard deviation (RSD) for all experimental values. I chose values for *k*s in the range of 10^3^ to 10^8^, and two isotope effect values in the classical range (1-6). With a RSD of 0.01, representing ideal experimental conditions, the modeled mean values are all within 10% of the true value, and the 90% confidence intervals contain the true value. Repeating this with other simulated values gives equally accurate results. The 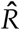statistic ^13,20^ measures the average divergence between MCMC chains during a simulation; in ideal data the value is 1.0 exactly, indicating that all the chains in the simulation have converged on the same posterior distribution. I have used 4 independent chains in each analysis. The 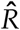 for all of the parameters in this investigation is less than 1.1, as prescribed by Ref. 13.

Increasing the RSD to 0.1, a much more realistic value, shows the model beginning to drift away from the true values and an increase in uncertainty. Nonetheless only two of the parameters is off by more than 50% – *k*_4_ and *k*_−3_. These two parameters are highly correlated, and in the absence of stronger prior information are likely to deviate from their simulated values. Notably, the ratio of *k*_4_ to *k*_−3_ is simulated as 0.5 and the fit shows 0.375, suggesting than an increase in prior information for either *k*_4_ or *k*_−3_ would greatly improve the estimate of both. While Toney validated his model with ideal datasets, his test data didn’t include experimental error and is analogous to my dataset with RSD = 0.01. The results of the modeling show that the current method is able to accurately determine rate constants under ideal conditions, and that experimental error begins to affect this at higher levels, as expected. I conclude from this that the model is structurally identifiable, with some parameters such as the intrinsic isotope effects determined with better precision than others.

### 3.2 Alanine Racemase

Table 3 shows the output of the Stan modeling for Alanine Racemase. We see that all of the parameters have converged well, as shown by the 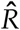 values lying close to unity. From the table, and from the graphs in Figure 11, the intrinsic KIEs (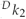 and ^*D*^ *k*_−3_) are in good agreement with the analysis from Ref. 1. The intrinsic KIEs for AR are especially well-defined, as shown by Figure 11. The prior and posterior distributions of both 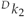 and ^*D*^ *k*_−3_ are shown, and the prior is much broader than the posterior, showing that the experimental data have been instumental in determining the mean and confidence intervals for the KIEs through the likelihood function. Others show some disagreement, especially the values of *k*_3_ and *k*_−2_ which differ by *>* 10-fold. The reason for this is not entirely clear, but is likely due to the effects of experimental uncertainty, as both in Ref. 1 and the present work the models are shown to give accurate results with ideal data. In the case of real world data, the difference between the models and algorithms becomes more important as error increases, as do the assumptions behind each. In every case where the present results and those of Ref. 1 disagree significantly (> 10-fold), the latter parameters show a great deal of uncertainty. In the case of AR, these are *k*_3_, *k*_−2_ and *k*_−3_. For *k*_3_ and *k*_−2_ we only have lower bounds in Ref. 1, and the SD of *k*_−3_ is 4 times the mean. In the present work, Stan is using the log of the joint likelihood function to estimate the shape and position of the posterior distribution, under the influence of a prior distribution; in Ref. 1 the program is trying to minimize a cost function (Eq. 1). Function minimization in the absence of a prior distribution can behave similarly to doing a Bayesian analysis with a Uniform prior on all parameters. In a case where the domain of each parameter is on the order of 10^12^, for a Bayesian analysis this gives us a prior where the parameter is nine times as likely to be in the range 10^11^ to 10^12^ as between 0 and 10^11^. This is part of the motivation for the use of the exponential distribution as a prior for all *k*s, to correct this bias towards larger numbers. In minimizing the function, if the experimental data are subject to error the parameters can vary freely over large ranges during the search, and converge to a wide range of values. This search is unbiased by a prior distribution, so might be preferred as long as the uncertainty in parameter estimates can be contained and there is little prior information. However in cases where parameters cannot be defined to within even an order of magnitude by function minimization, a Bayesian analysis such as the one shown here should be considered. I also note that the values that I estimate for all the parameters are consistent with the experimental data; the experimental means and theoretical mean values agree to within 1% in all cases. It may be that due to experimental uncertainty more than one set of parameters is consistent with the data (multimodality). In this case, an improvement of the experimental data or a more informative prior distribution might be necessary to resolve the problem.

**Table 1:** Experimental Values used to estimate Rate Constants.

|  | Alanine Racemase |  | Triosephosphate Isomerase |  |
| --- | --- | --- | --- | --- |
|  | Mean (SD) | Ref. | Mean (SD) | Ref. |
| $k_{cat,f}$ | 1740 (10) | 1 | 750 (50) | 1 |
| $k_{cat,r}$ | 1280 (12) | 1 | 8350 (350) | 1 |
| $K_{m,f}$ | 5.4 (0.1) | 1 | 1.35 (0.15) | 1 |
| $K_{m,r}$ | 4.0 (0.1) | 1 | 0.05 (0.01) | 1 |
| $K_{EZ}$ | $1.6 \times 10^{-4}$ ($0.4 \times 10^{-4}$) | 1 | $\leq 0.05$ | 1 |
| $\theta$ | 0.5 (0.1) | 1 | 3 (1) | 1 |
| slope | 0.015 (0.015) | 1 | 0.8 (0.1) | 1 |
| $^D k_{cat,f}$ | 1.5 (0.1) | 1 | 3.4 (0.1) | 1 |
| $^D k_{cat,r}$ | 1.4 (0.03) | 1 | 1.6 (0.1) | 1 |
| $^D \left( \frac{k_{cat}}{K_m} \right)_f$ | 1.6 (0.1) | 1 | 3.4 (0.1) | 1 |
| $^D \left( \frac{k_{cat}}{K_m} \right)_r$ | 1.3 (0.1) | 1 | 1.6 (0.1) | 1 |
| $K_{eq}$ | 1.0 (0.05) | 7 | 0.0035 (0.0001) | 8–10 |

**Table 2:** Statistical summary of the Stan output for the Simulated Data. 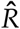 is the Gelman–Rubin statistic. ^20^ 90% CI is the 90% confidence interval for the posterior of each parameter. *RSD* is Relative Standard Deviation.

| Parameter | Sim. Value | Mean (RSD = 0.01) | 90% CI | $\hat{R}$ | Mean (RSD = 0.1) | 90% CI | $\hat{R}$ |
| --- | --- | --- | --- | --- | --- | --- | --- |
| $k_1$ | $2 \times 10^4$ | $1.97 \times 10^4$ | $(1.86 - 2.09) \times 10^4$ | 1.00 | $1.58 \times 10^4$ | $(1.39 - 1.80) \times 10^4$ | 1.00 |
| $k_2$ | $1 \times 10^3$ | $9.93 \times 10^2$ | $(9.56 - 10.3) \times 10^2$ | 1.00 | $7.51 \times 10^2$ | $(4.32 - 11.2) \times 10^2$ | 1.00 |
| $k_3$ | $1 \times 10^7$ | $1.00 \times 10^7$ | $(9.51 - 10.5) \times 10^6$ | 1.00 | $1.40 \times 10^7$ | $(7.07 - 27.2) \times 10^6$ | 1.00 |
| $k_4$ | $1 \times 10^4$ | $1.06 \times 10^4$ | $(8.61 - 13.1) \times 10^3$ | 1.00 | **$3.45 \times 10^6$** | **$(9.22 - 75.4) \times 10^5$** | 1.00 |
| $k_{-1}$ | $5 \times 10^3$ | $4.94 \times 10^3$ | $(4.68 - 5.23) \times 10^3$ | 1.00 | $3.99 \times 10^3$ | $(3.91 - 4.10) \times 10^3$ | 1.00 |
| $k_{-2}$ | $1 \times 10^8$ | $9.81 \times 10^7$ | $(9.43 - 10.2) \times 10^7$ | 1.00 | $6.69 \times 10^7$ | $(3.81 - 10.1) \times 10^7$ | 1.00 |
| $k_{-3}$ | $2 \times 10^4$ | $2.15 \times 10^4$ | $(1.73 - 2.66) \times 10^4$ | 1.00 | **$9.19 \times 10^6$** | **$(2.05 - 20.4) \times 10^6$** | 1.00 |
| $k_{-4}$ | $1 \times 10^5$ | $9.92 \times 10^4$ | $(9.67 - 10.2) \times 10^4$ | 1.00 | $8.96 \times 10^4$ | $(7.22 - 11.3) \times 10^4$ | 1.00 |
| $^Dk_{-3}$ | 3.5 | 3.48 | (3.44 – 3.52) | 1.00 | 3.26 | (2.86 – 3.66) | 1.00 |
| $^Dk_{-3}$ | 1.5 | 1.53 | (1.48 – 1.58) | 1.00 | 1.95 | (1.35 – 2.81) | 1.00 |

**Table 3:** Statistical summary of the Stan output for the AR Data. 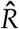 is the Gelman–Rubin statistic. ^20^ 90% CI is the 90% confidence interval for the posterior of each parameter.

| Parameter | Mean | Median | 90% CI | $\hat{R}$ | Num. Opt. (Toney, 2013) |
| --- | --- | --- | --- | --- | --- |
| $k_1$ (mM<sup>-1</sup>s<sup>-1</sup>) | $1.4 \times 10^4$ | $1.3 \times 10^4$ | $(7.2 \times 10^3, 2.6 \times 10^4)$ | 1.00 | $> 10^5$ |
| $k_2$ (s<sup>-1</sup>) | $2.8 \times 10^3$ | $2.8 \times 10^3$ | $(2.5 \times 10^3, 3.0 \times 10^3)$ | 1.00 | 2600(200) |
| $k_3$ (s<sup>-1</sup>) | $8.3 \times 10^6$ | $8.2 \times 10^6$ | $(6.3 \times 10^6, 1.1 \times 10^7)$ | 1.00 | $1.4 \times 10^7(0.3 \times 10^7)$ |
| $k_4$ (s<sup>-1</sup>) | $9.0 \times 10^5$ | $7.2 \times 10^5$ | $(2.5 \times 10^5, 2.1 \times 10^6)$ | 1.00 | $9 \times 10^6(8 \times 10^7)$ |
| $k_{-1}$ (s<sup>-1</sup>) | $7.5 \times 10^4$ | $6.7 \times 10^4$ | $(3.7 \times 10^4, 1.4 \times 10^5)$ | 1.00 | $> 10^6$ |
| $k_{-2}$ (s<sup>-1</sup>) | $4.7 \times 10^6$ | $4.6 \times 10^6$ | $(6.7 \times 10^5, 5.9 \times 10^6)$ | 1.00 | $6.8 \times 10^6(0.7 \times 10^6)$ |
| $k_{-3}$ (s<sup>-1</sup>) | $3.8 \times 10^3$ | $3.7 \times 10^3$ | $(3.2 \times 10^3, 4.5 \times 10^3)$ | 1.00 | 4000(700) |
| $k_{-4}$ (mM<sup>-1</sup>s<sup>-1</sup>) | $2.1 \times 10^2$ | $1.7 \times 10^2$ | $(5.6 \times 10^1, 4.9 \times 10^2)$ | 1.00 | $2 \times 10^4(2 \times 10^5)$ |
| $^Dk_2$ | 1.56 | 1.56 | (1.44, 1.67) | 1.00 | 1.55(0.11) |
| $^Dk_{-3}$ | 1.41 | 1.41 | (1.36, 1.46) | 1.00 | 1.35(0.09) |

**Table 4:** Statistical summary of the Stan output for the TIM Data. 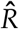 is the Gelman–Rubin statistic. ^20^ 90% CI is the 90% confidence interval for the posterior of each parameter.

| Parameter | Mean | Median | 90% CI | $\hat{R}$ | Num. Opt. (Ref. 1) |
| --- | --- | --- | --- | --- | --- |
| $k_1$ ($\text{mM}^{-1}\text{s}^{-1}$) | $1.4 \times 10^4$ | $1.3 \times 10^4$ | $(9.7 \times 10^3, 2.2 \times 10^4)$ | 1.01 | 8400(1500) |
| $k_2$ ($\text{s}^{-1}$) | $4.4 \times 10^3$ | $4.2 \times 10^3$ | $(2.9 \times 10^3, 6.3 \times 10^3)$ | 1.00 | 4000(1600) |
| $k_3$ ($\text{s}^{-1}$) | $9.7 \times 10^4$ | $8.3 \times 10^4$ | $(4.1 \times 10^4, 1.9 \times 10^5)$ | 1.00 | $> 10^9$ |
| $k_4$ ($\text{s}^{-1}$) | $1.3 \times 10^4$ | $1.3 \times 10^4$ | $(9.5 \times 10^3, 1.8 \times 10^4)$ | 1.00 | $1.2 \times 10^4(0.3 \times 10^4)$ |
| $k_{-1}$ ($\text{s}^{-1}$) | $2.0 \times 10^4$ | $1.9 \times 10^4$ | $(1.4 \times 10^4, 3.1 \times 10^5)$ | 1.00 | $1.1 \times 10^4(0.2 \times 10^4)$ |
| $k_{-2}$ ($\text{s}^{-1}$) | $7.9 \times 10^4$ | $7.2 \times 10^6$ | $(4.0 \times 10^4, 1.4 \times 10^5)$ | 1.00 | $> 10^8$ |
| $k_{-3}$ ($\text{s}^{-1}$) | $5.3 \times 10^4$ | $4.8 \times 10^4$ | $(2.8 \times 10^4, 9.8 \times 10^4)$ | 1.00 | $5 \times 10^5(20 \times 10^5)$ |
| $k_{-4}$ ($\text{mM}^{-1}\text{s}^{-1}$) | $4.0 \times 10^4$ | $3.8 \times 10^4$ | $(2.2 \times 10^4, 6.3 \times 10^4)$ | 1.00 | $2.3 \times 10^5(0.3 \times 10^5)$ |
| $^Dk_2$ | 3.57 | 3.57 | (3.45, 3.71) | 1.00 | 3.56(0.08) |
| $^Dk_{-3}$ | 2.64 | 2.61 | (2.10, 3.31) | 1.00 | 3.3(0.5) |

### 3.3 Triosephosphate Isomerase

For TIM, similar issues arise as for AR. There is an assumption that the uncertainties in each parameter are distributed normally. Looking at the estimates for the TIM dataset, Ref. 1 estimated *k*_−3_ as 5 × 10^5^ with a SD (20 × 10^5^) ; if you assume normalcy that would mean a full 40% of the confidence interval lies below zero, where it is impossible for the value to be. Likewise for *k*_−2_, the optimization results give wide ranges for the parameter confidence interval.

Nickbarg and Knowles ^9^ also calculated the ratios of the forward and reverse rate constants for yeast TIM. Table 5 shows a comparison of the results from the current study compared with Ref. 1 and Nickbarg and Knowles (1988). There is agreement between the ratios of the forward and reverse constants for the first and fourth steps (*k*_1_/*k*_−1_ and *k*_4_/*k*_−4_). The current work and Nickbarg and Knowles (1988) give essentially identical estimates for *k*_2_/*k*_−2_ and *k*_3_/*k*_−3_, with Ref. 1 differing by ≈ 10^3^ in both cases. At stake is the question of whether the complex of TIM with the enediol intermediate (EZ, in Scheme 1) is significantly higher in energy than the other enzyme forms. Higher energy would destabilize EZ, leading to higher *k*_−2_ and *k*_3_ values and would therefore lead to *k*_2_/*k*_−2_ approaching zero and *k*_3_/*k*_−3_ much greater than one. While it is not my intention to wade into this debate, the results presented here are not consistent with a high-energy intermediate.

**Table 5:** Comparison of forward and reverse rate-constant ratios from the present work, Nickbarg and Knowles (1988), and Toney (2013). Results in the two rightmost columns are given as Mean (SD).

|  | Mean | 90% CI | Nickbarg and Knowles (1988) | Toney (2013) |
| --- | --- | --- | --- | --- |
| $k_1/k_{-1}$ | 0.7 | (0.6, 0.9) | 0.6 (0.3) | 0.8 (0.2) |
| $k_2/k_{-2}$ | 0.05 | (0.02, 0.09) | 0.1 (0.5) | $< 4 \times 10^{-5}$ |
| $k_3/k_{-3}$ | 1.8 | (0.8, 3.6) | 2 (10) | $> 2000$ |
| $k_4/k_{-4}$ | 0.06 | (0.04, 0.09) | 0.04 (0.02) | 0.05 (0.01) |

### 3.4 Free Energy Profiles for TIM and AR

Starting with the MCMC draws for the eight rate constants, we can construct a series of FEPs and overlay them in an attempt to map the most likely profile for each of the enzyme. In Figures 8 and 9, I have randomly chosen 500 of the 4000 draws for each MCMC run, and calculated the FEP for that particular draw using the Eyring equation with *κ* = 1 and *T* = 298*K*. The FEPs show a great deal of overlap when plotted on top of each other, and species with uncertain energies show up as a series of lines, as in the EZ intermediate of AR (Figure 8). In Figures 8 and 9, I have also calculated the averrage energy of each species, shown as a red horizontal line.

**Figure 8:**
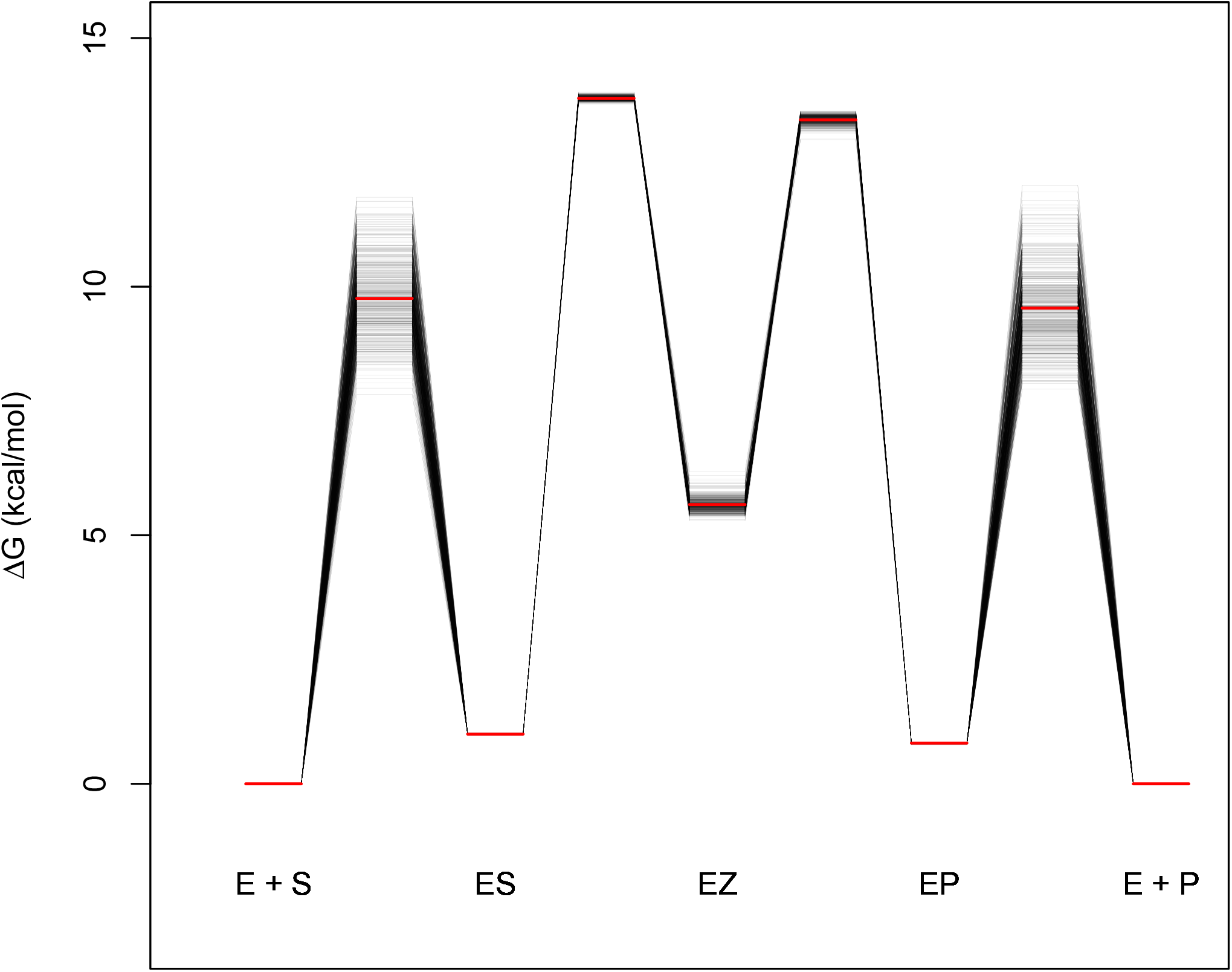
The Free Energy Profile for the AR reaction, relative to the starting substrate. The red lines are the means, for a given species.

**Figure 9:**
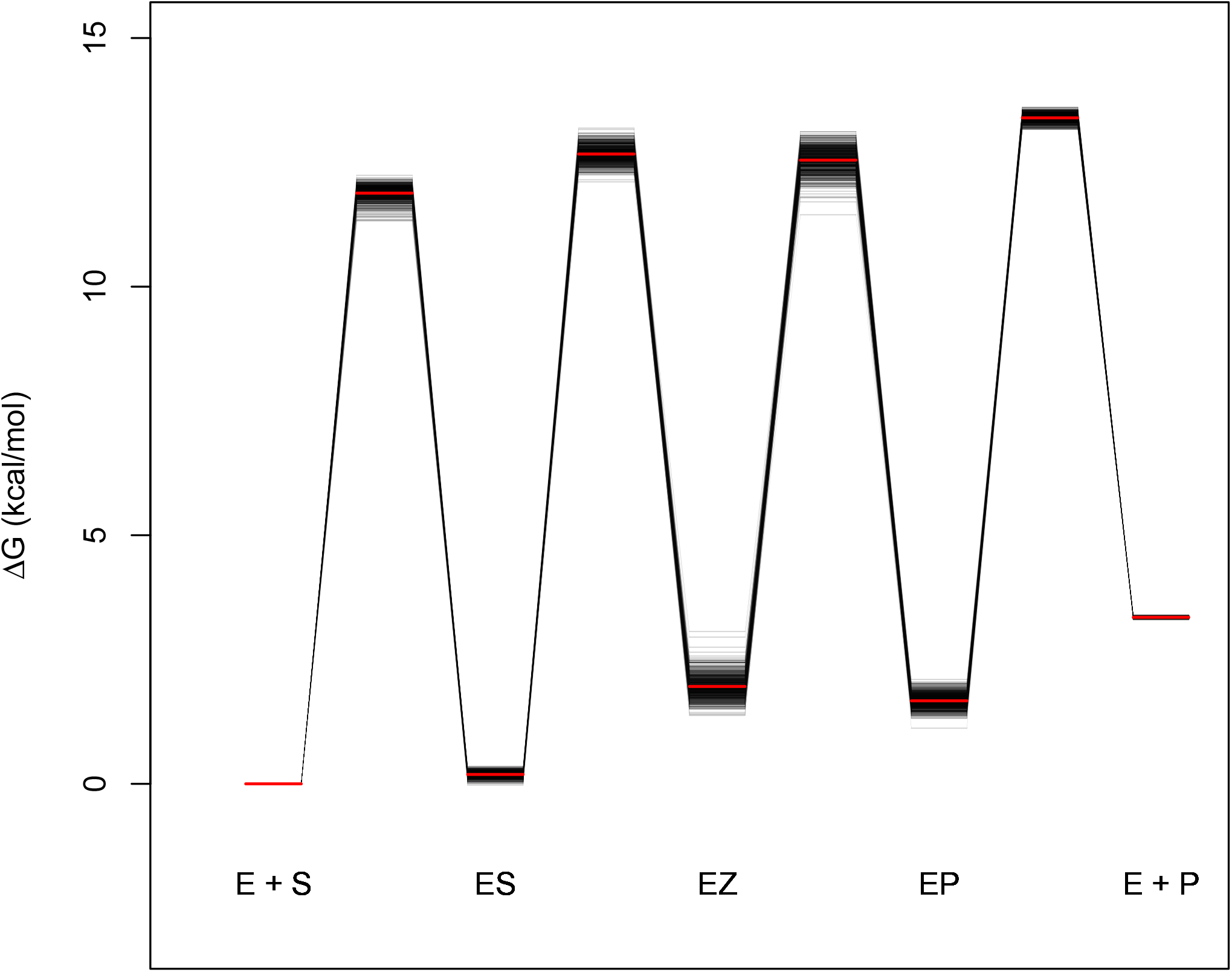
The Free Energy Profile for the TIM reaction, relative to the starting substrate. The red lines are the means, for a given species.

## 4. Methods

### 4.1 Incorporation of Experimental Error

The data ^1^ is expressed as mean 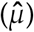 and standard deviation 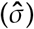 as is commonly done in biochemical studies. The data are incorporated into the model as follows:

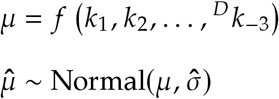

This is represented in Stan as follows, for *k*_*cat, f*_ (ignoring all other data values):

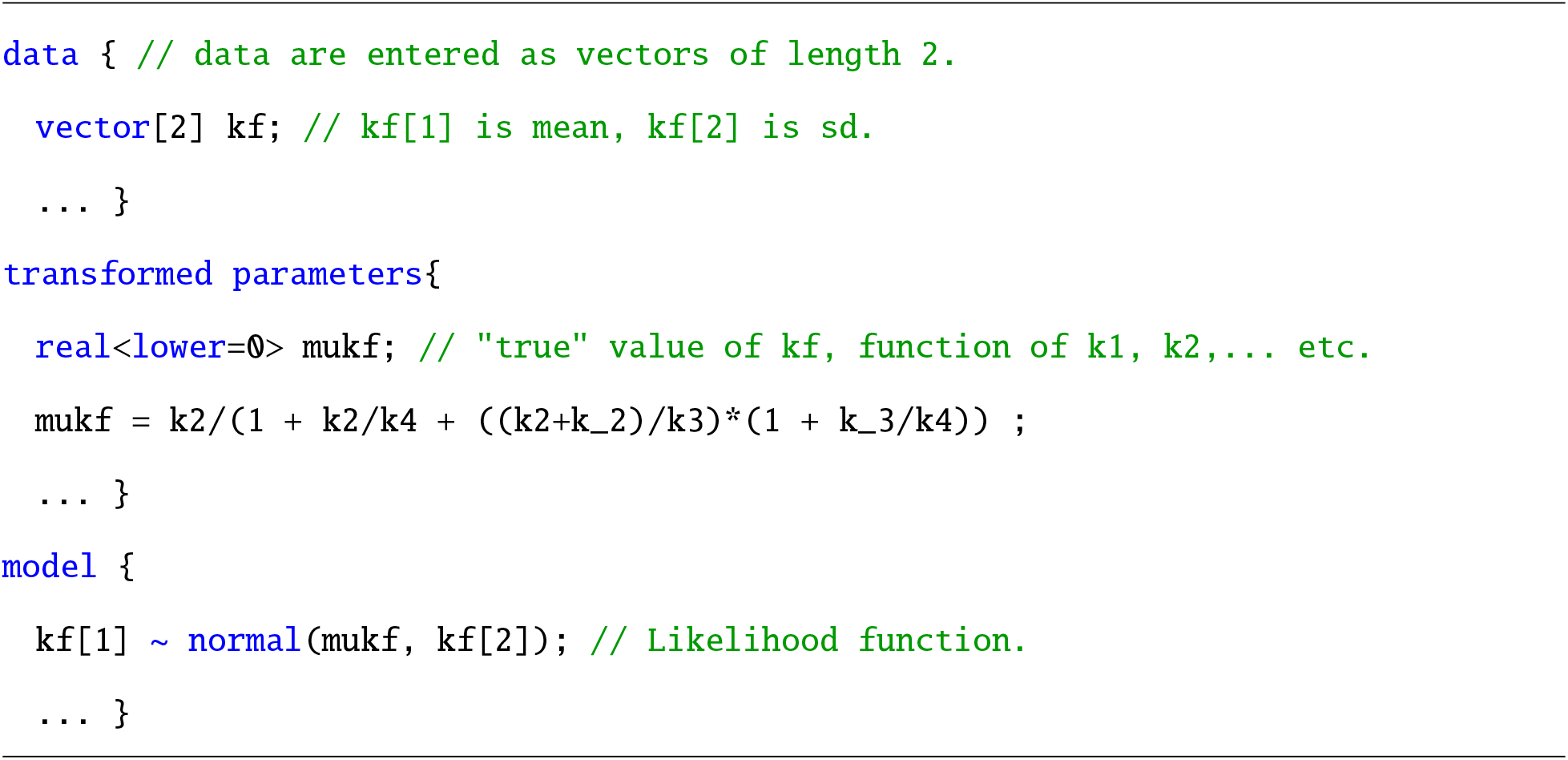

Where the true value of the experimentally-determined parameter (e.g. *k*_*cat*_) is assumed to be drawn from a distribution with mean *µ*, and the standard deviation is set equal to the experimentally-determined uncertainty in the value. The model will then incorporate the mean value 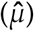 as data, and estimate a true value for it as well (*µ*), based on the global fitting. While one could use a distribution other than Normal to model the error, most published results use models that assume Normally-distributed error so when incorporating results from others it is important to follow this assumption. Ideally, the model would incorporate the raw data instead and proceed from there to estimates of the rate constants; however, the raw data is not available, as is often the case with biochemical data. Nonetheless it is still possible to get estimates of the rate constants from published results.

### 4.2 MCMC Analysis

To estimate the posterior distribution of each of the eight parameters, I used cmdstanr 0.3.0 running under R 3.5, which is built on cmdstan 2.26.1. ^21^ For each analysis, 5000 iterations of the sampler were run on 4 parallel chains. The first 4000 of each were ‘warm-up’ samples, in which the parameters and step sizes are tuned by Stan’s NUTS algorithm. This value, higher than the default value of 1000 warm-up samples, was necessary to ensure that the sampling distribution was stable, but had minimal effect on the runtime of the program. Runtimes on MacOS using a 3 GHz quad-core processor and 4 parallel chains ranged from 2 - 60s, without diagnostic errors after sampling. Figures were generated using ggplot2 and the bayesplot package, except Figure 10 which was plotted with gnuplot.

**Figure 10:**
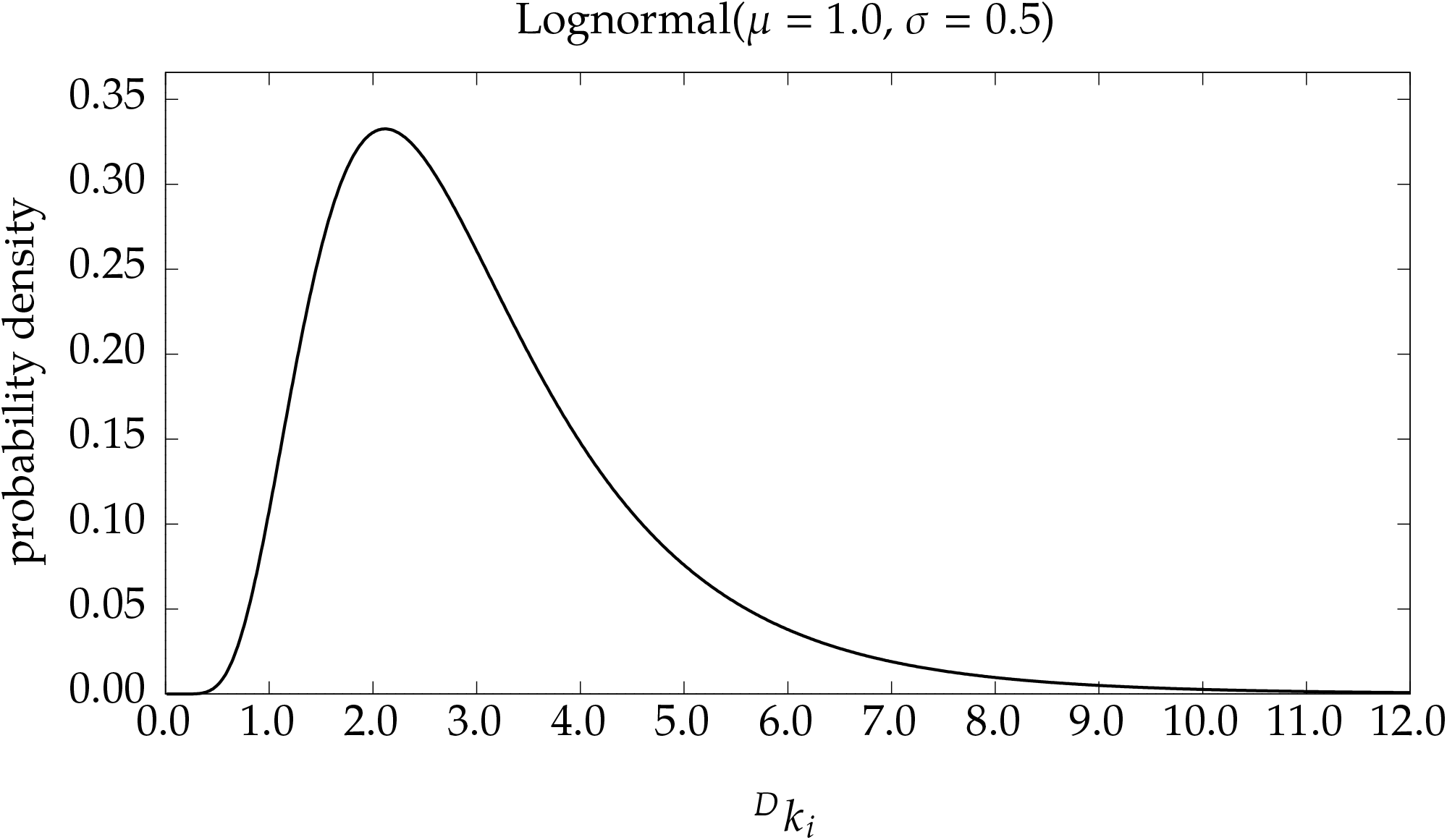
The prior distribution used for intrinsic KIEs (^*D*^ *k*_*i*_)

**Figure 11:**
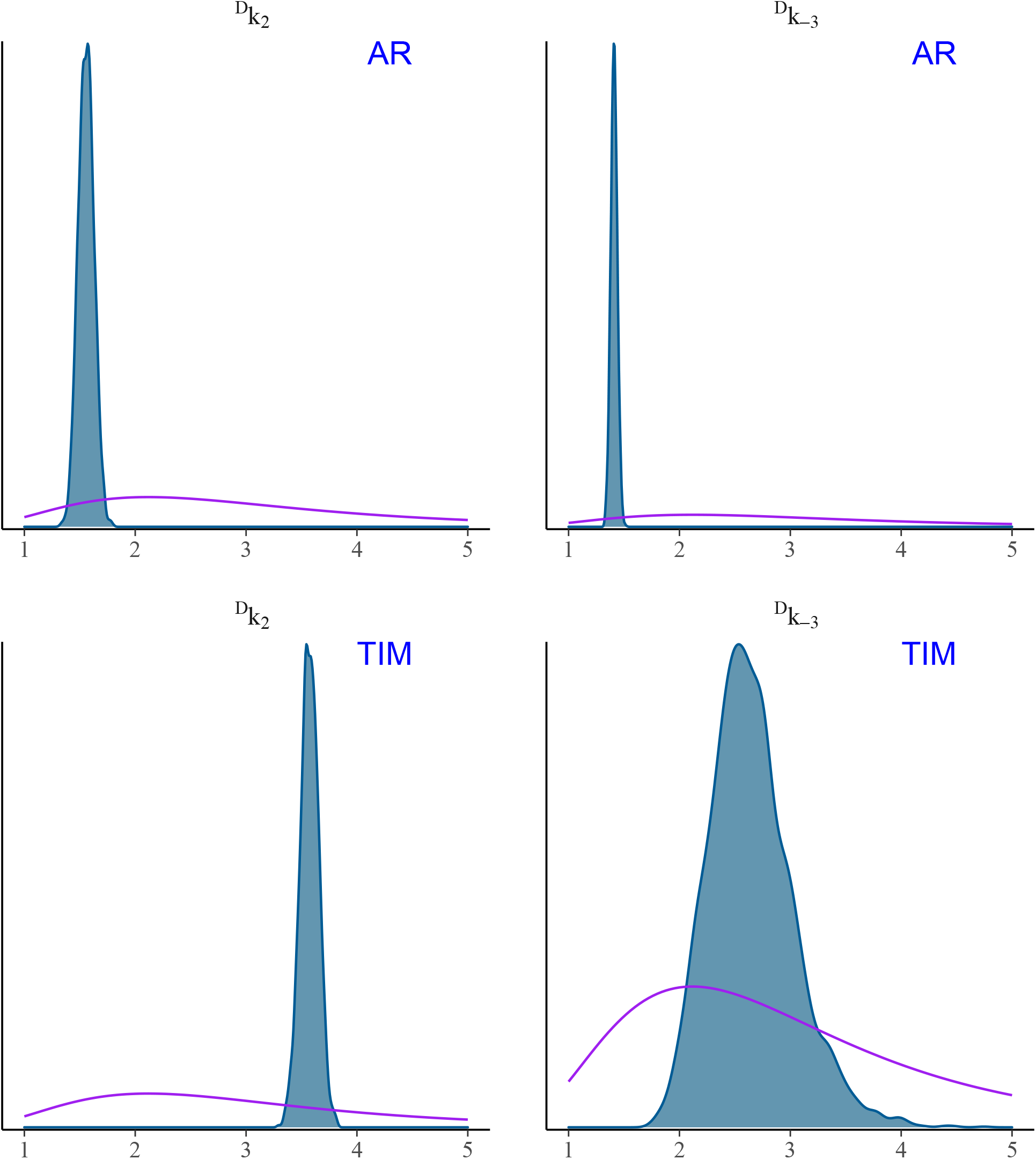
Posterior and prior distributions for the intrinsic KIEs (^*D*^ *k*_*i*_) of TIM and AR. The prior distribution is shown in purple, and the posterior is shown in blue, filled.

## Acknowledgement

This work was partially supported by a Summer Research Grant from Dominican University of California.

## Supporting Information Available

The following files are available free of charge.

- simulate.stan: Stan model file
- enrg.R: R script file to process Toney (2013) data.
- simulate.R: R script file to simulate and process data.

## References

(1) Toney, M. D. Common enzymological experiments allow free energy profile determination. Biochemistry 2013, 52, 5952–65.

(2) Cornish-Bowden, A. Fundamentals of Enzyme Kinetics, 3rd ed.; Portland Press Limited: London, 2004.

(3) James, F.; Roos, M. Minuit - a system for function minimization and analysis of the parameter errors and correlations. Computer Physics Communications 1975, 10, 343–367.

(4) Gallant, A. R. Nonlinear Regression. The American Statistician 1975, 29, 73–81.

(5) McNeish, D. On Using Bayesian Methods to Address Small Sample Problems. Structural Equation Modeling: A Multidisciplinary Journal 2016, 23, 750–773.

(6) Betancourt, M. A Conceptual Introduction to Hamiltonian Monte Carlo. 1701.02434. 2018.

(7) Goldberg, R. N.; Tewari, Y. B. Thermodynamics of Enzyme-Catalyzed Reactions: Part 5. Isomerases and Ligases. Journal of Physical and Chemical Reference Data 1995, 24, 1765–1801.

(8) Albery, W. J.; Knowles, J. R. Free-energy profile of the reaction catalyzed by triosephosphate isomerase. Biochemistry 1976, 15, 5627–31.

(9) Nickbarg, E. B.; Knowles, J. R. Triosephosphate isomerase: energetics of the reaction catalyzed by the yeast enzyme expressed in Escherichia coli. Biochemistry 1988, 27, 5939–47.

(10) Sampson, N. S.; Knowles, J. R. Segmental motion in catalysis: investigation of a hydrogen bond critical for loop closure in the reaction of triosephosphate isomerase. Biochemistry 1992, 31, 8488–94.

(11) Cleland, W. W. An analysis of Haldane Relationships. Methods Enzymol 1982, 87, 366–369.

(12) Fersht, A. Structure and mechanism in protein science: a guide to enzyme catalysis and protein folding; W.H. Freeman: New York, 1999.

(13) Gelman, A.; Carlin, J. B.; Stern, H. S.; Dunson, D. B.; Vehtari, A.; Rubin, D. B. Bayesian Data Analysis, third edition ed.; Chapman & Hall/CRC texts in statistical science; 2013.

(14) Clauset, A.; Shalizi, C. R.; Newman, M. E. J. Power-Law Distributions in Empirical Data. SIAM Review 2009, 51, 661–703.

(15) Mitzenmacher, M. A Brief History of Generative Models for Power Law and Lognormal Distributions. Internet Mathematics 2004, 1, 226–251.

(16) Klinman, J. P. The role of tunneling in enzyme catalysis of C-H activation. Biochim Biophys Acta 2006, 1757, 981–7.

(17) Hu, S.; Sharma, S. C.; Scouras, A. D.; Soudackov, A. V.; Carr, C. A. M.; Hammes-Schiffer, S.; Alber, T.; Klinman, J. P. Extremely elevated room-temperature kinetic isotope effects quantify the critical role of barrier width in enzymatic C-H activation. J Am Chem Soc 2014, 136, 8157– 8160.

(18) Dalziel, K. Physical significance of Michaelis constants. Nature 1962, 196, 1203–5.

(19) Bellman, R.; Åström, K. On structural identifiability. Mathematical Biosciences 1970, 7, 329–339.

(20) Gelman, A.; Rubin, D. B. Inference from Iterative Simulation Using Multiple Sequences. Statistical Science 1992, 7, 457–472.

(21) Stan Development Team, Stan Modeling Language Users Guide and Reference Manual, version 2.27. mc-stan.org 2019,.

