## Supplementary material for "A Bayesian Analysis of Steady–State Enzyme Data leads to Estimates of Rate Constants and Uncertainties in a Multi-Step Reaction, along with Free Energy Profiles^†^": enrg.R

```
library("cmdstanr")
library("bayesplot")
library("ggplot2")
set.seed(1)
## compile the Stan model
enrg <- cmdstan_model("simulate.stan")

## Data for AR, in form data = c(mean,sd)
dataAR <- list(
  kf = c(1740, 10),
  kr = c(1280 ,12 ),
  Kmf = c(5.4 ,0.1 ),
  Kmr = c(4.0 , 0.1 ),
  Dkf = c(1.5 ,0.1 ),
  Dkr = c(1.4 ,0.03 ),
  DKmr = c(1.5,0.1 ),
  DKmf = c(1.6 ,0.1 ),
  slope = c( 0.015, 0.015),
  theta = c(0.5 , 0.1),
  kez = c(1.6e-4 , 4e-5),
  Keq = c(1.0, .001)
)
## sample the posterior
outptAR <- enrg$sample(data = dataAR, seed = 1234, iter_warmup =  4e3, iter_sampling = 1e3, chains = 4, parallel_chains = 4, max_treedepth = 20, adapt_delta = 0.85 )
outptAR.df <- outptAR$summary()
ARdraws <- outptAR$draws()
## Data for TIM, in form data = c(mean,sd)
dataTIM <- list(
  kf = c(750, 50),
  kr = c(8350 ,350 ),
  Kmf = c(1.35 ,0.15 ),
  Kmr = c(0.05 , 0.01 ),
  Dkf = c(3.4 ,0.1 ),
  DKmr = c(1.6,0.1 ),
  DKmf = c(3.4 ,0.1 ),
  Dkr = c(1.6 ,0.1 ),
  slope = c( 0.8, 0.1),
  theta = c(3 , 1),
  kez = c(0.025 , 0.025),
  Keq = c(0.0035, 0.0001)
  #Keq = c(0.045, 0.01)
)
## sample the posterior
outptTIM <- enrg$sample(data = dataTIM, seed = 1234, chains = 4, iter_warmup =  4e3, iter_sampling = 1e3, parallel_chains = 4, max_treedepth = 20, adapt_delta = 0.85 )
outptTIM.df <- outptTIM$summary()
TIMdraws <- outptTIM$draws()
#######
#####graphing the results
#####labels for graphs
label <- as_labeller(x = c(
  # old_name = "new name"
  lp__   = "test" ,
  K1    = 'K[1]', 
  k1    = 'k[1]',  
  k2   = 'k[2]' ,   
  k3   = 'k[3]' ,   
  k4  = 'k[4]' ,    
  k_1  = 'k[-1]' ,   
  k_2  = 'k[-2]' ,   
  k_3   = 'k[-3]' ,  
  k_4  = 'k[-4]' ,
  K4  = 'K[4]' ,
  Dk2  = 'phantom(0)^D*k[2]',
  Dk_3  = 'phantom(0)^D*k[-3]' ,
  mukf  = "test" ,
  mukr  = "test" ,
  muKmf = "test" ,
  muKmr = "test" ,
  muDkf = "test" ,
  muDkr  = "test" ,
  muDKmf = "test" ,
  muDKmr = "test" ,
  mutheta = "test" ,
  muslope= "test" ,
  mukez = "test" ,
  dG1  = "test" ,
  dG2 = "test" ,
  dG3  = "test" ,
  dG4  = "test"  
),  default = label_parsed)

#plots for AR
theme_set(theme_default())

traceplotAR <- mcmc_trace(ARdraws, pars = c("k1","k2","k3","k4","k_1","k_2","k_3","k_4"), facet_args = list(labeller = label))
ggsave(traceplotAR, filename = 'traceplotAR.pdf')

theme_set(theme_default() +  xaxis_text(on = FALSE) +yaxis_text(on = FALSE))

pairplotAR <- mcmc_pairs(ARdraws, pars = c("k1","k2","k3","k4","k_1","k_2","k_3","k_4", "Dk2", "Dk_3"), 
                         diag_fun = "dens", off_diag_fun = "hex", grid_args = list(labeller = label) )
ggsave(pairplotAR, filename = 'pairplotAR.pdf')

theme_set(theme_default())
traceplotTIM <- mcmc_trace(TIMdraws, pars = c("k1","k2","k3","k4","k_1","k_2","k_3","k_4"), facet_args = list(labeller = label))
ggsave(traceplotTIM, filename = 'traceplotTIM.pdf')

theme_set(theme_default() +  xaxis_text(on = FALSE) + yaxis_text(on = FALSE))

pairplotTIM <- mcmc_pairs(TIMdraws, pars = c("k1","k2","k3","k4","k_1","k_2","k_3","k_4", "Dk2", "Dk_3"),
                          diag_fun = "dens", off_diag_fun = "hex", grid_args = list(labeller = label))
ggsave(pairplotTIM, filename = 'pairplotTIM.pdf')
theme_set(theme_default())


theme_set(theme_default() + theme(aspect.ratio = 1))# + coord_fixed())
#pdf(file = "TIM-Dkprior.pdf")
lnorm_prior <-
  overlay_function(
    fun = dlnorm,
    args = list(meanlog = 1, sdlog = 0.5),
    color = "purple",
    size = 0.5
  )
priorplotTIM <- mcmc_dens(TIMdraws, pars = c("Dk2","Dk_3"), facet_args = list(labeller = label)) + 
  annotate("text", x=4.5, y=Inf, label= "TIM", colour = "blue", size = 5, vjust=1, hjust=1) + lnorm_prior + xlim(1,5) #+ ylim(0,5.5)
#print(priorplotTIM)
#dev.off()

#####AR graph
#pdf(file = "AR-Dkprior.pdf")


priorplotAR <- mcmc_dens(ARdraws, pars = c("Dk2","Dk_3"), facet_args = list(labeller = label)) + 
  annotate("text", x=4.5, y=Inf, label= "AR", colour = "blue", size = 5, vjust=1, hjust=1) + lnorm_prior + xlim(1,5) #+ ylim(0,15) 
    
# print(priorplotAR)
# dev.off()

priorplotKIEs <- bayesplot_grid(priorplotAR,priorplotTIM)
ggsave(priorplotKIEs, filename = 'Dkprior.pdf')

theme_set(theme_default())
outptTIM$save_output_files(dir = "./TIM/")
outptAR$save_output_files(dir = "./AR/")


eyring <- function(k, T){
  h <- 6.6261e-34 #J.s
  kB <- 1.3806e-23 #J/K
  R <- 1.9872e-3 # kcal/(K.mol)
  dG <- -R*T*log((k*h)/(kB*T))
  return(dG)
}

pdf(file = "AR-EnergyPlot.pdf")
ARdraws.df <- posterior::as_draws_df(ARdraws)
ARz <- ARdraws.df[ , c("k1", "k_1", "k2", "k_2","k3", "k_3", "k4", "k_4")]
AR_ENERGY <- plot(NULL, xlim=c(0,18), ylim=c(-3,15), ylab=expression(paste(Delta,"G"," (kcal/mol)")), xlab="",xaxt="n")
AR_N <- 500
dGs <- matrix(0.0, nrow = AR_N, ncol = 9)

drwz <- ARz[sample(nrow(ARz), AR_N), ]
for (i in 2:9){
  l <- (-1)^(i)
  #drwz = sample(ARz[[i-1]], 1000)
  dGs[,i] = dGs[,(i-1)] + l*eyring(drwz[[i-1]],298)
  segments(x0=rep(1, AR_N), y0=rep(0, AR_N), x1=rep(2.0, AR_N), y1=rep(0, AR_N), lw = 0.1)
  segments(x0=rep(2.0*i-1, AR_N), y0=dGs[,i], x1=rep(i*2.0, AR_N), y1=dGs[,i], lw = 0.1, col='#00000030')
  segments(x0=rep(2.0*(i-1.0), AR_N), y0=dGs[,(i-1)], x1=rep(i*2.0-1.0, AR_N), y1=dGs[,i], lw = 0.1, col='#00000050')
  text(c(2., 6., 10., 14., 18.)-0.5, rep(-2, 5), c("E + S", "ES", "EZ", "EP", "E + P"))
}
mudG <- c(mean(dGs[,1]),mean(dGs[,2]),mean(dGs[,3]),mean(dGs[,4]),mean(dGs[,5]),mean(dGs[,6]),mean(dGs[,7]),mean(dGs[,8]),mean(dGs[,9]))
segments(x0 = (2*1:9 -1), y0 = mudG, x1 = (2*1:9 ), y1 = mudG, lw = 1.5, col = "red")
print(AR_ENERGY)
dev.off()


pdf(file = "TIM-EnergyPlot.pdf")
TIMdraws.df <- posterior::as_draws_df(TIMdraws)
TIMz <- TIMdraws.df[ , c("k1", "k_1", "k2", "k_2","k3", "k_3", "k4", "k_4")]
TIM_ENERGY <- plot(NULL, xlim=c(0,18), ylim=c(-3,15), ylab=expression(paste(Delta,"G"," (kcal/mol)")), xlab="",xaxt="n")

TIM_N <- 500
dGs <- matrix(0.0, nrow = TIM_N, ncol = 9)
drwz <- TIMz[sample(nrow(TIMz), TIM_N), ]
for (i in 2:9){
  l <- (-1)^(i)
  #drwz = sample(ARz[[i-1]], 1000)
  dGs[,i] = dGs[,(i-1)] + l*eyring(drwz[[i-1]],298)
  segments(x0=rep(1, TIM_N), y0=rep(0, TIM_N), x1=rep(2.0, TIM_N), y1=rep(0, TIM_N), lw = 0.5)
  segments(x0=rep(2.0*i-1, TIM_N), y0=dGs[,i], x1=rep(i*2.0, TIM_N), y1=dGs[,i], lw = .5, col='#00000030')
  segments(x0=rep(2.0*(i-1.0), TIM_N), y0=dGs[,(i-1)], x1=rep(i*2.0-1.0, TIM_N), y1=dGs[,i], lw = 0.1, col='#00000030')
  text(c(2., 6., 10., 14., 18.)-0.5, rep(-2, 5), c("E + S", "ES", "EZ", "EP", "E + P"))
  
}
mudG <- c(mean(dGs[,1]),mean(dGs[,2]),mean(dGs[,3]),mean(dGs[,4]),mean(dGs[,5]),mean(dGs[,6]),mean(dGs[,7]),mean(dGs[,8]),mean(dGs[,9]))
segments(x0 = (2*1:9 -1), y0 = mudG, x1 = (2*1:9 ), y1 = mudG, lw = 1.5, col = "red")
print(TIM_ENERGY)
dev.off()
```
