## Supplementary material for "A Bayesian Analysis of Steady–State Enzyme Data leads to Estimates of Rate Constants and Uncertainties in a Multi-Step Reaction, along with Free Energy Profiles^†^": simulate.R

```
library("cmdstanr")
library("bayesplot")
library("ggplot2")

func_data <- function(datvec) {
k1 <- datvec[1] 
k2 <- datvec[2] 
k3 <- datvec[3] 
k4 <- datvec[4]
k_1 <- datvec[5] 
k_2 <- datvec[6] 
k_3 <- datvec[7] 
k_4 <- datvec[8] 
Dk2 <- datvec[9]
Dk_3 <- datvec[10]
tkf <- k2/(1 + k2/k4 + ((k2+k_2)/k3)*(1 + k_3/k4)) 
ttheta <- (k_2*(1+k_3/k4))/(k3*(1+k2/k_1))
tkr <- k_3/(1 + k_3/k_1 + ((k3+k_3)/k_2)*(1 + k2/k_1)) 
tkez <- 1/(k_2/k2 + k3/k_3 +1)
tslope <- ( k_1*k_2*k_3 + k2*k3*k4)/(k_1*k_2*k_3 + k2*k3*k4 + k_1*k3*k4 + k_1*k_2*k4)
tKmf <- (k_1/k1)*(1 + k2/k_1 +(k_2/k3)*(1+k_3/k4))/(1 + k2/k4 + ((k2 + k_2)/k3)*(1+k_3/k4))
tKmr <- (k4/k_4)*(1 + k_3/k4 +(k3/k_2)*(1+k2/k_1))/(1 + k_3/k_1 + ((k3 + k_3)/k_2)*(1+k2/k_1))
tDkf <- ( Dk2 + (k2/k3)*(1+k_3/k4) + k2/k4 + Dk2*(k_2/k3)*(1+k_3/k4)) / ( 1 + (k2/k3)*(1+k_3/k4) + k2/k4 + (k_2/k3)*(1+k_3/k4))
tDkr <- ( Dk_3 + (k_3/k_2)*(1+k2/k_1) + k_3/k_1 + Dk_3*(k3/k_2)*(1+k2/k_1))/( 1 + (k_3/k_2)*(1+k2/k_1) + k_3/k_1 + (k3/k_2)*(1+k2/k_1))
tDKmf <- ( Dk2 + k2/k_1 + Dk2*(k_2/k3)*(1+k_3/k4)) / ( 1 + (k2/k_1) + (k_2/k3)*(1+k_3/k4))
tDKmr <- ( Dk_3 + (k_3/k4) + Dk_3*(k3/k_2)*(1+k2/k_1))/(  1 + (k_3/k4) + (k3/k_2)*(1+k2/k_1) )
tKeq <- (k1*k2*k3*k4)/(k_1*k_2*k_3*k_4)
return(c(tkf, tkr, tkez, tslope, tKmf, tKmr, tDkf,tDkr, tDKmf,tDKmr,ttheta, tKeq ))
}
#ks <- c(2e4,1e3, 1e7,1e4, 5e3, 1e8,2e4,1e5, 3.5, 1.5)
ks <- c(1e4,1e4, 1e4,1e4, 1e4,1e4, 1e4,1e4, 2.5, 2.5)
test_mean <- func_data(ks)
test_matrix  <- matrix(nrow = 12, ncol = 6)
set.seed(3321)
for (i in 1:4){
  test_matrix[,i] <- test_mean * rnorm(12, mean = 1.0, sd  = 0.01)
}
for (j in 1:12){
  test_matrix[j,5] <- mean(test_matrix[j,1:4])
  test_matrix[j,6] <- sd(test_matrix[j,1:4])
}

#
dataSIM <- list(
  kf = c(test_matrix[1,5], test_matrix[1,6]),
  kr = c(test_matrix[2,5], test_matrix[2,6]),
  kez = c(test_matrix[3,5], test_matrix[3,6]),
  slope = c(test_matrix[4,5], test_matrix[4,6]),
  Kmf = c(test_matrix[5,5], test_matrix[5,6]),
  Kmr = c(test_matrix[6,5], test_matrix[6,6]),
  Dkf = c(test_matrix[7,5], test_matrix[7,6]),
  Dkr = c(test_matrix[8,5], test_matrix[8,6]),
  DKmf = c(test_matrix[9,5], test_matrix[9,6]),
  DKmr = c(test_matrix[10,5], test_matrix[10,6]),
  theta = c(test_matrix[11,5], test_matrix[11,6]),
  Keq = c(test_matrix[12,5], test_matrix[12,6])
)
init <- function(){
  list(
  k1 = runif(1, 1e3, 1e6), 
  k2 = runif(1, 1e3, 1e6), 
  k3 = runif(1, 1e3, 1e6), 
  k4 = runif(1, 1e3, 1e6),
  k_1 = runif(1, 1e3, 1e6), 
  k_2 = runif(1, 1e3, 1e6), 
  k_3 = runif(1, 1e3, 1e6), 
  k_4 = runif(1, 1e3, 1e6), 
  Dk2 = runif(1, 1, 6),
  Dk_3 = runif(1, 1, 6))}
enrg <- cmdstan_model("simulate.stan")
outptSIM <- enrg$sample(data = dataSIM, seed = 1234, iter_warmup =  4e3, iter_sampling = 1e3, chains = 4, parallel_chains = 4, max_treedepth = 20, adapt_delta = .95 , init = init )
outptSIM.df <- outptSIM$summary()
SIMdraws <- outptSIM$draws()

theme_set(theme_default() +  xaxis_text(on = FALSE) + yaxis_text(on = FALSE))

pairplotSIM <- mcmc_pairs(SIMdraws, pars = c("k1","k2","k3","k4","k_1","k_2","k_3","k_4", "Dk2", "Dk_3"),
                          diag_fun = "dens", off_diag_fun = "hex", grid_args = list(labeller = label))
ggsave(pairplotSIM, filename = 'pairplotSIM.pdf')
theme_set(theme_default())

traceplotSIM <- mcmc_trace(SIMdraws, pars = c("k1","k2","k3","k4","k_1","k_2","k_3","k_4"), facet_args = list(labeller = label))
ggsave(traceplotSIM, filename = 'traceplotSIM.pdf')
theme_set(theme_default())
```
